# Malaria parasite resistance to azithromycin is not readily transmitted by mosquitoes

**DOI:** 10.1101/2023.11.10.566666

**Authors:** Hayley D. Buchanan, Robyn McConville, Lee M. Yeoh, Michael F. Duffy, Justin A. Boddey, Geoffrey I. McFadden, Christopher D. Goodman

**Affiliations:** School of BioSciences, University of Melbourne, Parkville, 3010, Vic, Australia; Walter and Eliza Hall Institute of Medical Research, Parkville, 3052, Vic, Australia; Department of Medical Biology, University of Melbourne, Parkville, 3010, Vic, Australia; Bio21 Institute, Parkville, 3052, Vic, Australia; Department of Microbiology and Immunology, School of Biomedical Sciences, University of Melbourne, Parkville, 3010, Vic, Australia

## Abstract

Antimalarials are now used in combination with partner drugs to stem parasite drug resistance. Partners are often older, safe, cheap drugs, but resistance is already circulating for many, which raises the risk of selecting for multidrug resistance. If the partner drug(s) could be refractory to the spread of resistance, better resistance control could be implemented. We tested whether resistance to the antibiotic azithromycin, which kills malaria parasites by perturbing prokaryote-like protein synthesis in the apicoplast (relict plastid), had fitness costs to the spread of parasites via mosquitoes where parasites are not under drug pressure. Azithromycin resistance mutations in both rodent and human malaria parasites had a negative impact on the ability of resistant parasites to transmit from one vertebrate host to another via mosquitoes. Azithromycin resistance will therefore be less likely to spread geographically, making it an attractive option as a perennial partner compound to protect appropriate frontline antimalarials.

## INTRODUCTION

Drug resistance is a major challenge to controlling, let alone eradicating, malaria^1^. Resistant parasites emerge, spread geographically and eventually erode drug efficacy. A Sisyphean cycle of using-then-losing malaria drugs has repeated itself at least four times, starting in the 1950s with chloroquine and currently playing out with artemisinin, resistance to which has now spread through the Greater Mekong Delta ^2–6^ and emerged in Africa ^6–8^. Clearly, we need to do things differently to gain more control of malaria drug resistance and stop relying on one promising new drug to replace the previous one that we have set up for failure.

Combination therapies have slowed the spread of artemisinin resistance, and any new frontline drug will require protection by some form of partner compound(s). Triple and even quadruple combinations are now being considered ^9, 10^, but where do we turn for the partner drugs? Older, inexpensive drugs with acceptable safety profiles are one option, but resistance is already circulating for many, raising the risk of selecting for multidrug resistance ^11^. Ideally, the partner compounds should not only be safe, cheap, compatible, and effective but also be refractory to the spread of resistance. This would prolong the life of the frontline compound whilst also minimising the risk of selecting for multidrug resistance.

Atovaquone is an antimalarial refractory to the spread of resistance ^12, 13^. Atovaquone resistance is unable to be transmitted because resistant parasites with mutations in the mitochondrially-encoded cytochrome *b* suffer fitness deficits during the mosquito phase of their life cycle where they rely on higher rates of mitochondrial electron transport ^13, 14^. In search of further resistance-refractory drugs, we turned to other drug/target systems with similar life cycle constraints. We hypothesised that changes to the parasite fitness landscape when switching from vertebrate to mosquito hosts might also occur in other metabolic pathways that are more active in the insect and/or liver phases of the life cycle. An obvious candidate is the apicoplast, a relict plastid of malaria parasites that is substantially more active in mosquito and liver stages ^15–17^ and for which a wide selection of safe, cheap inhibitors is known ^18–20^.

The apicoplast is a non-photosynthetic plastid homologous to the chloroplasts of plants and algae ^21^. Apicoplasts have their own DNA and express proteins encoded by their genome using housekeeping machinery within the organelle ^22^. Apicoplasts, like their chloroplast relatives, arose by endosymbiosis of a photosynthetic cyanobacterium ^23^, and contain their own DNA replication, RNA transcription, and protein translation and modification machinery, which is bacterial in nature and sensitive to a raft of antibacterials ^24^. For this reason, various antibacterials, including azithromycin, are also parasiticidal ^19, 20, 24–29^.

Azithromycin is a broad-spectrum macrolide antibiotic that blocks apicoplast translation by binding to the peptide exit tunnel of the 50S subunit of the apicoplast ribosome ^30^. Like many other drugs targeting the apicoplast, azithromycin has a key drawback as a therapeutic—its lethal effect is delayed until the second intraerythrocytic after treatment. This “delayed death” response results primarily from apicoplast metabolic fatigue that prevents parasite feeding during blood stage, resulting in slow starvation ^31^. Despite the delayed drug effects, azithromycin is used as monotherapy to treat uncomplicated malaria and in combination with chloroquine or artesunate ^27–29^. There are calls to increase the use of azithromycin as an antimalarial for combinations ^32^, and this push is strengthened by the recent discovery that azithromycin has two targets in malaria parasites: one in the apicoplast, and a second that inhibits parasite invasion of the red blood cell ^33, 34^. Azithromycin is safe for infants and pregnant women and has a very long half-life in the body, all of which make it attractive as a partner compound ^32^.

Investigations into the role of the apicoplast in malaria parasites across the two-host life cycle show that the organelle is essential at every stage ^35^, but the roles of the apicoplast wax and wane across the life cycle ^36^. In the blood stage the apicoplast has only two roles: to manufacture the five-carbon isoprenoid precursor isopentenyl diphosphate ^18, 37^, and to synthesise coenzyme A (CoA) ^38^. By contrast, in the mosquito oocyst stage the apicoplast also has to make fatty acids ^15^ and heme ^39–41^, although these metabolic requirements are also parasite species-specific ^36^. The apicoplast thus has different roles—and hence demands—in the two hosts, which implies that selection constraints on apicoplast efficiency will differ across the life cycle, much as they do with mitochondrial electron transport ^13, 14, 42^.

We hypothesised that malaria parasites resistant to azithromycin will experience differential selection across the parasite life cycle and incur a deferred fitness cost that restricts their ability to develop in mosquitoes and/or liver stages. Thus, apicoplast mutations allowing parasites to survive under drug pressure in the blood stage should be sub-optimal in mosquito and/or liver stages, when drug pressure is released. To test our hypothesis, we generated azithromycin resistant rodent (*Plasmodium berghei*) and human (*P. falciparum*) malaria parasites and assayed their ability to transmit from vertebrate-to-vertebrate via mosquitoes.

## RESULTS

### Generation of resistance to azithromycin in vivo in P. berghei and in vitro in P. falciparum

Azithromycin resistance has been reported in malaria parasites ^30, 33^, but to assay resistance transmissibility, we generated new azithromycin resistant parasites in genetic backgrounds able to be transmitted in the laboratory. In *P. berghei* ANKA, several cycles of treatment→recrudescence→treatment with azithromycin at the ED_95_ (60 mg/kg q.d.; Figure 1A and B, Figure S1) or ED_99_ (70 mg/kg q.d.; Figure 1A and B, Figure S1) yielded three, independently derived lines of *P. berghei* azithromycin resistant parasites: PbAZMR_G95D_1, PbAZMR_G95D_2 (both selected on a 70 mg/kg q.d. drug regimen), and PbAZMR_S89L (selected on 60 mg/kg q.d.). The delayed-death kinetics of azithromycin make *ex vivo* drug trials impossible for *P. berghei* so resistance phenotypes were confirmed *in vivo*. Stable resistance was observed at the maximal allowable azithromycin dose of twice the ED_99_ dose (140 mg/kg q.d.) (Figure 1C), but animal ethics constraints prevented treatment at higher doses and, therefore, precluded ED_50_ determinations for these resistant strains. The slow growth rate of PbAZMR_S89L (Figure S2) prevented *in vivo* confirmation of resistance after multiple attempts failed to produce high enough blood stage parasitemia and exflagellations.

**Figure 1.**
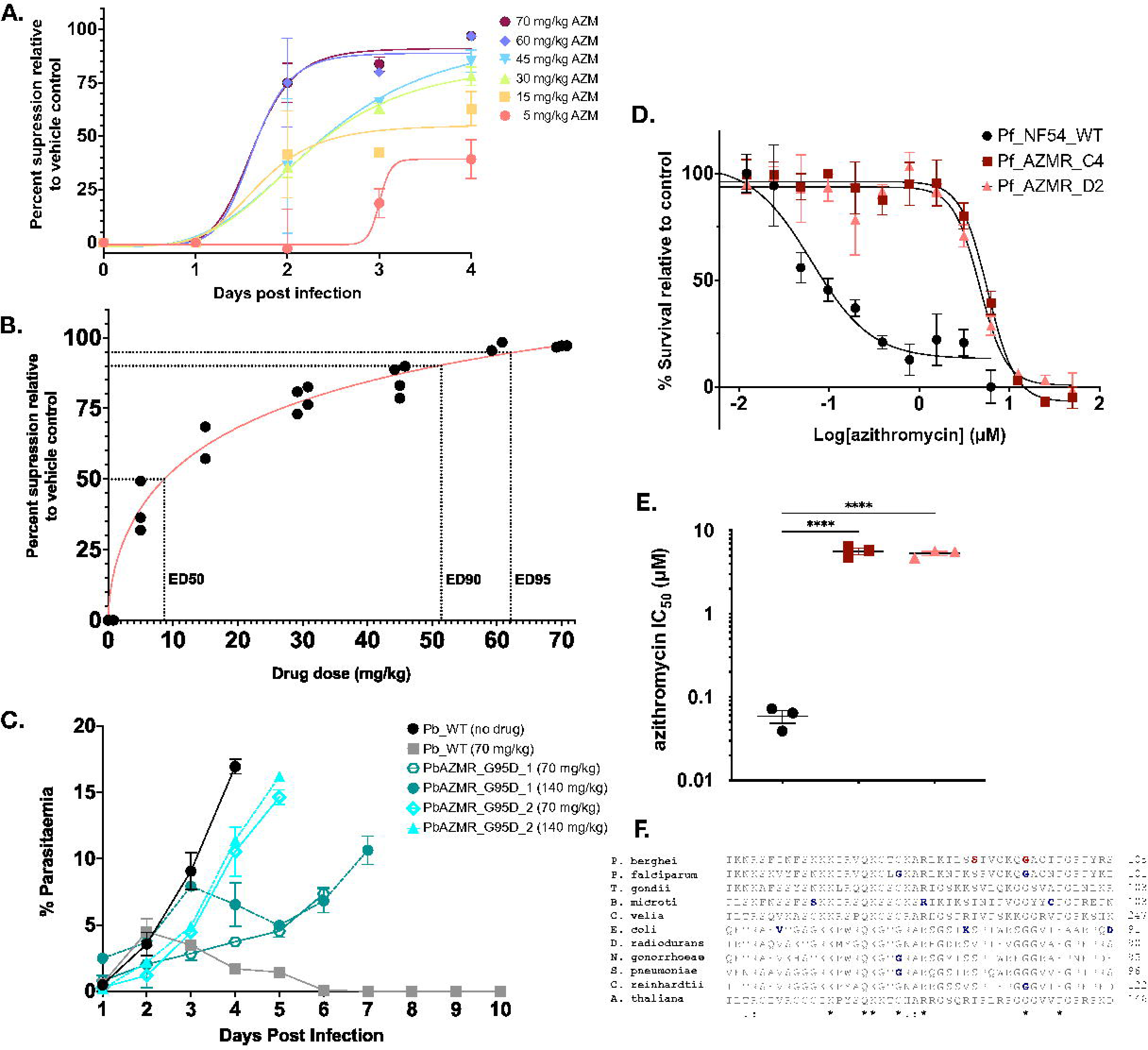
Summary drug assays performed *in vivo* with *P. berghei* and *in vitro* with *P. falciparum*. A.) Peters’ four-day suppressive test *in vivo* with *P. berghei*. Percent suppression of a variable dosing range used to determine both the appropriate clearance dose and potential resistance-generating dose. Parasitaemias of individual mice in each dose cohort were normalised to the mean vehicle control-treated parasitaemia at each time point, and the mean percent-suppression value plotted (error bars represent ±SD). A non-linear regression curve with a variable slope was then fitted to the data points for each dose cohort; B.) End-point (day 4) data for the Peters’ four-day suppressive test. Parasitaemias of individual mice in each dose cohort were normalised to the mean vehicle control-treated parasitaemia at day 4 and each individual value plotted. A non-linear regression curve with a variable slope was then fitted to the data points. Where n<4, mice were euthanised prior to the experimental end point for humane reasons; C.) *In vivo* drug assay of PbAZMR_G95D_1 (teal) and PbAZMR_G95D_2 (light blue) grown with a full four-day 70 mg/kg q.d. (solid lines, n=2 per genotype) or 140 mg/kg q.d. (dashed lines, n=2 per genotype) drug course versus WT with the vehicle (no drug; black; n=2) and a full four-day 70 mg/kg q.d. drug course (grey; n=2). Treatment was started on day 1. (Values indicate mean parasitaemia, and error bars represent the SEM); D.) Dose response curves of Pf_AZMR_C4 and Pf_AZMR_D2 clones (both with G76V mutation in *P. falciparum* 50S Rpl4) compared to Pf_WT. Data presented are the mean of three technical replicates and representative of three independent drug trials (error bars represent ±SD); E.) Comparison plot of IC50s of *P. falciparum* azithromycin resistant clones versus WT. Plotted are the mean IC50 values from three independent drug trials. (error bars represent the SEM, **** = P<0.0001, ordinary one-way ANOVA, Dunnett’s multiple comparisons test); F.) Alignment of region of Rpl4 associated with azithromycin of erythromycin resistance in bacteria and plastids of parasites, algae and plants. Mutations identified in this study are depicted in red and mutations selected from the literature in blue. Included organisms: P. berghei (*Plasmodium berghei*); P. falciparum (*Plasmodium falciparum*); T. gondii (*Toxoplasma gondii*); B. microti (*Babesia microti*); C. velia (*Chromera velia*), E.coli (*Escherichia coli*); D. radiodurans (*Deinococcus radiodurans*); N. gonorrhoeae (*Neisseria gonorrhoeae*); S. pneumoniae (*Streptococcus pneumoniae*); C. reinhardtii (*Chlamydomonas reinhardtii*);A.thaliana (*Arabidopsis thaliana*). Asterisks denote invariant residues, colons depect conserved residues groups with strongly similar properties, and periods denote semi-conserved residues groups with similar properties. (AZM = azithromycin; ED = effective dose)

We selected for azithromycin resistance *in vitro* in the *P. falciparum* mosquito-transmissible strain NF54 as previously described ^30^. The IC_50_ of azithromycin in a 120-hour, delayed-death assay for the parental line was 0.059 ± 0.017 μM (Figure 1D and E). Two independent cultures of a clonal line of NF54, and one of a clonal line of the non-transmissible 3D7 strain, were grown in the presence of 300 nM azithromycin for 196 hours and then maintained in drug-free culture for 24-days, at which time parasites recrudesced. Resistance to azithromycin in both *P. falciparum* strains was achieved in a single treatment cycle. Time to recrudescence was similar for control parasites selected for resistance to pyrimethamine or atovaquone. An increase of drug to 600 nM azithromycin to azithromycin selected cultures did not inhibit growth (Figure 1D). Resistance to azithromycin at IC_50_s ∼100-fold greater than the parental line was maintained in all lines following cloning (Figure 1D and E).

### Azithromycin resistance in P. berghei and P. falciparum is apparently conferred by mutations in Rpl4

It is well established that azithromycin targets apicoplast translation in malaria parasites, specifically the 50S ribosomal subunit ^20, 30^. Moreover, reported resistance to azithromycin in malaria parasites is conferred by apicoplast-encoded mutations ^30, 33, 43^. Whole genome sequencing of several clones of each of our independently generated *P. berghei* and *P. falciparum* azithromycin resistant lines revealed mutations in a conserved region of the apicoplast-encoded 50S ribosomal protein L4 (Rpl4; PBANKA_API00430 in *P. berghei* and PF3D7_API01300 in *P. falciparum* respectively; Figure 1F). Two different point mutations in Rpl4, S89L and G95D, were found in *P. berghei* (Figure 1F). Another mutation, G76V, was found in Rpl4 of azithromycin resistant *P. falciparum* (Figure 1F). These Rpl4 point mutations were observed in the absence of any other identifiably relevant point mutations, indels, or copy number variations.

### Azithromycin resistant P. berghei parasites are severely impaired during mosquito stage development

To test the sexual viability of our *P. berghei* azithromycin resistant strains we quantified the numbers of exflagellating males before each infection. There were no significant differences in the number of exflagellations seen in PbAZMR_G95D_1, PbAZMR_G95D_2, and the Pb_WT (Figure 2A). It is unclear if the reduction in PbAZMR_S89L parasites was a specific effect or resulted from the poor blood stage growth of this mutant (Figure S2). Furthermore, this blood stage growth defect limited us to only two independent experiments, so statistical analysis was not robust.

**Figure 2.**
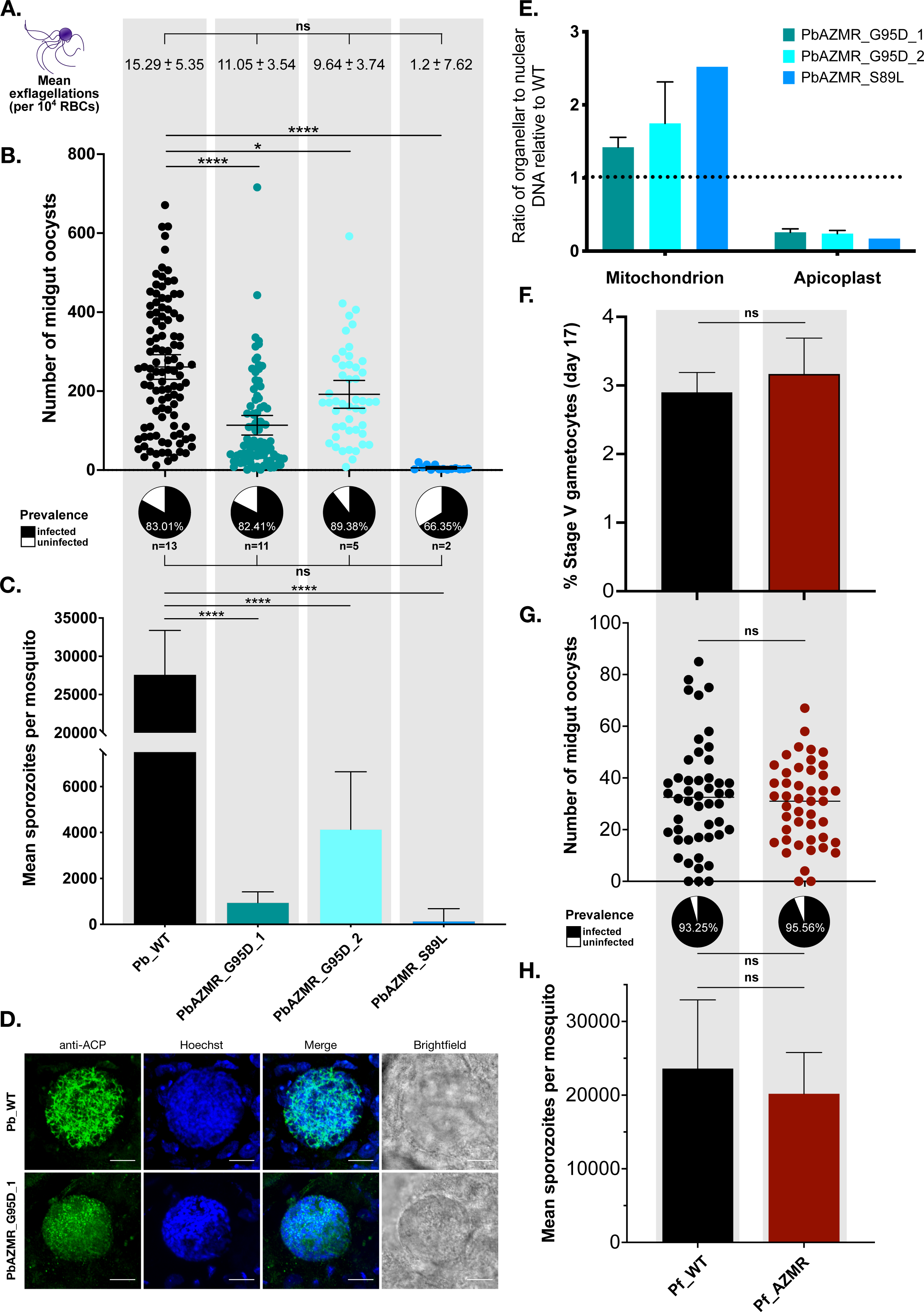
Mosquito infection data for both *P. berghei* and *P. falciparum* azithromycin resistant parasites. A.) Exflagellations were counted per 1×10^4^ red blood cells (RBCs) and are shown as the mean ±95%CI, with no significant difference between strains (ordinary one-way ANOVA with Dunnett’s multiple comparisons; WT [n=17], PbAZMR_G95D_1 [n=20], PbAZMR_G95D_2 [n=10], PbAZMR_S89L [n=2], where ‘n’ is the number of independent biological replicates). At 9-to-14 days post blood feed, a minimum of 10 midguts were dissected for each ‘n’ independent infection (WT [n=13], PbAZMR_G95D_1 [n=11], PbAZMR_G95D_2 [n=5], PbAZMR_S89L [n=2]) and assessed for infection prevalence (pie charts below, percentage indicates the mean; ns= not significant by way of ordinary one-way ANOVA). Oocysts of infected midguts were counted and pooled, and non-zero values were plotted (error bars represent the mean ±95%CI). All azithromycin resistant strains had significantly fewer oocysts than WT (*=P<0.05; ****=P<0.0001; ordinary one-way ANOVA, Dunnett’s multiple comparisons test); B.) Mean number of salivary gland sporozoites per mosquito. Between days 20- and 26-post blood feed, a minimum of 12 mosquitoes (WT) or 20 mosquitoes (PbAZMR_G95D_1, PbAZMR_G95D_2, and PbAZMR_S89L) were dissected. All azithromycin resistant strains produce fewer salivary gland sporozoites than WT (****=P<0.0001, Welch’s t-test; error bar represents +95% CI). Results are from ‘n’ independent mosquito infections: WT (n=21), PbAZMR_G95D_1 (n=14), PbAZMR_G95D_2 (n=10), PbAZMR_S89L (n=2); C.) Quantitative PCR on genomic DNA isolated from midguts infected with both *P. berghei* azithromycin resistant parasites and WT strains. Copy numbers of genes encoded in each organelle were first normalised to nuclear genes, then expressed as a ratio to WT parasites. There is a clear reduction in apicoplast genome copy number in PbAZMR_G95D_1 (n=4), PbAZMR_G95D_2 (n=3) and PbAZMR_S89L (n=1), where n=number of biological replicates. Error bars represent the SEM; D.) Immunofluorescence assay on midgut oocysts, 14 days post blood feed. Green represents labelling of the *P. berghei* apicoplast, with anti-ACP antibodies. Hoechst is staining DNA. PbAZMR_G95D_1 oocysts consistently show dispersed ACP staining (bottom panel), while WT oocysts show a branched ACP staining (top panel). Images are maximum projections (except brightfield, which shows the largest single slice). Scale bar=10μm; E.) Quantification of development of blood-feed-ready stage V gametocytes produced by Pf_WT and Pf_AZMR parasites. No significant difference is seen (Unpaired t-test, two-tailed P value); F.) At 7 days post blood feed, a minimum of 10 midguts were dissected for each ‘n’ independent infection (Pf_WT [n=3], Pf_AZMR [n=3]) and assessed for infection prevalence (pie charts below, percentage indicates the mean; ns= not significant by way of Unpaired t-test, two-tailed P value). No significant difference was observed between the number of midgut oocysts in mosquitoes infected with either Pf_WT or Pf_AZMR parasites (Unpaired t-test, two-tailed P value); G.) At day 17 post blood feed, a minimum of 12 mosquitoes were dissected per genotype. No significant difference was observed between the number of salivary gland sporozoites in mosquitoes infected with either Pf_WT or Pf_AZMR parasites (Unpaired t-test, two-tailed P value).

We next infected female *Anopheles stephensi* mosquitoes with our azithromycin resistant *P. berghei* lines and Pb_WT control parasites by allowing the mosquitoes to bite infected mice. In a minimum of five independent mosquito infections with PbAZMR_G95D_1, PbAZMR_G95D_2, and Pb_WT strains, there was no significant difference in the prevalence of mosquito infection, but the resistant lines developed significantly fewer oocysts with means of 114 ± 12.4, 191± 17.5, and 261 ± 15.8 oocysts per midgut respectively (Figure 2B). The development impairment in PbAZMR_S89L parasites during mosquito midgut stages likely reflects a mix of specific mosquito stage effects and very slow blood stage growth producing fewer sexual stages (Figure 2A) and therefore fewer oocysts.

At 20-26 days post blood feed, salivary glands were dissected from mosquitoes infected with PbAZMR_G95D_1, PbAZMR_G95D_2, PbAZMR_S89L, and Pb_WT. Again, all our azithromycin resistant lines produced significantly fewer salivary gland sporozoites than Pb_WT (Figure 2C). The reduction in sporozoite numbers was markedly greater than would be expected from the lower number of oocysts observed, and suggests a defect in development during the oocyst stage.

To visualise oocyst stage development in our azithromycin resistant *P. berghei* strains, we performed an immunofluorescence assay (IFA) on whole, infected mosquito midguts (Figure 2D). *P. berghei* azithromycin resistant oocysts had abnormal, dispersed staining with the apicoplast-directed antibody to anti-acyl carrier protein (ACP), whereas Pb_WT oocysts showed the expected branched apicoplast pattern (Figure 2D). ACP is encoded in the parasite nucleus and targeted to the apicoplast ^44, 45^ and continues to be produced even if the apicoplast is not intact. The dispersed ACP staining seen in our PbAZMR oocysts suggests that the apicoplast structure has been disrupted, and ACP is likely accumulating in vesicles destined for the non-existent or deformed apicoplast. Although IFA of whole midguts is informative, it is not quantitative, partly because the epithelial and basal lamina cells of the *A. stephensi* midgut and the *P. berghei* oocyst wall are significant barriers to consistent reagent entry and prevent consistent antibody staining of oocysts.

To quantify the developmental defect in the apicoplasts of azithromycin resistant *P. berghei* oocysts we measured their organllear DNA by quantitative PCR (qPCR), which revealed a loss of between 74.3% and 82.8% of apicoplast genomes in the *P. berghei* azithromycin resistant strains compared to Pb_WT (Figure 2E). Loss of the apicoplast genome confirms the microcsopic observations that azithromycin resistance-conferring mutations in apicoplast-encoded Rpl4 disrupt apicoplast development during mosquito stages in *P. berghei*. The concomitant increase in mitochondrial genome copy number in all the azithromycin resistant strains compared to Pb_WT (Figure 2E) was not expected but could be due to abberent nuclear development (Figure 2D) and disruption of the timing of genome replication, perturbing normalisation of organelle genome number to nuclear gene numbers.

### Azithromycin resistant P. falciparum parasites develop normally in mosquitoes

To determine if azithromycin resistance mutations impeded *P. falciparum* mosquito stage development, we selected a single clone (Pf_AZMR_C4, hereafter Pf_AZMR) to conduct downstream analyses. We compared the development of Pf_AZMR, and the parent line *P. falciparum* NF54 wildtype (Pf_WT) in female *A. stephensi* mosquitoes following standard membrane feeding. Both Pf_AZMR and Pf_WT parasite lines produced equivalent numbers of mature stage V gametocytes (Figure 2F). Similarly, both Pf_AZMR and Pf_WT parasites produced equivalent numbers of midgut oocysts at day 7 post blood feed, with an equivalent infection prevalence (Figure 2G). Furthermore, Pf_AZMR and Pf_WT parasites had no significant difference in the number of salivary gland sporozoites at day 17 post blood feed (Figure 2H). Thus, Pf_AZMR, despite having a point mutation in Rpl4 near those that severely disrupt mosquito infectivity of Rpl4 mutants in *P. berghei*, is apparently uninhibited in its ability to infect mosquitoes.

### Azithromycin resistant P. berghei sporozoites lose their apicoplast

Immunofluorescence assays of apicoplast morphology in PbAZMR_G95D_1 and PbAZMR_G95D_2 sporozoites revealed the absence of a defined apicoplast, with only diffuse ACP staining visible throughout the length of the sporozoite (Figure 3A). This constrasts with Pb_WT sporozoites that have a canonical, punctate apicoplast structure adjacent the nucleus as expected (Figure 3A). The diffuse ACP staining seen in azithromycin resistant sporozoites (Figure 3A) is reminiscent of the dispersed apicoplast marker labelling in oocysts of azithromycin resistant parasites and may again indicate futile ACP trafficking. To quantify this aberration, we used IFA microscopy to assess >100 sporozoites from each parasite strain, across three independent mosquito infections (Figure 3B). Only 12.6% of PbAZMR_G95D_1 sporozoites, and 18.8% of PbAZMR_G95D_2 sporozoites exhibited intact apicoplasts, whereas 93% of Pb_WT sporozoites had clearly identifiable, punctate apicoplasts (Figure 3B). Apparently, the disruption of apicoplast development in oocysts that results from the azithromycin resistance-conferring Rpl4 mutations reduces apicoplast integrity during *P. berghei* mosquito stage development, culminating in the production of sporozoites that predominantly lack an apicoplast.

**Figure 3.**
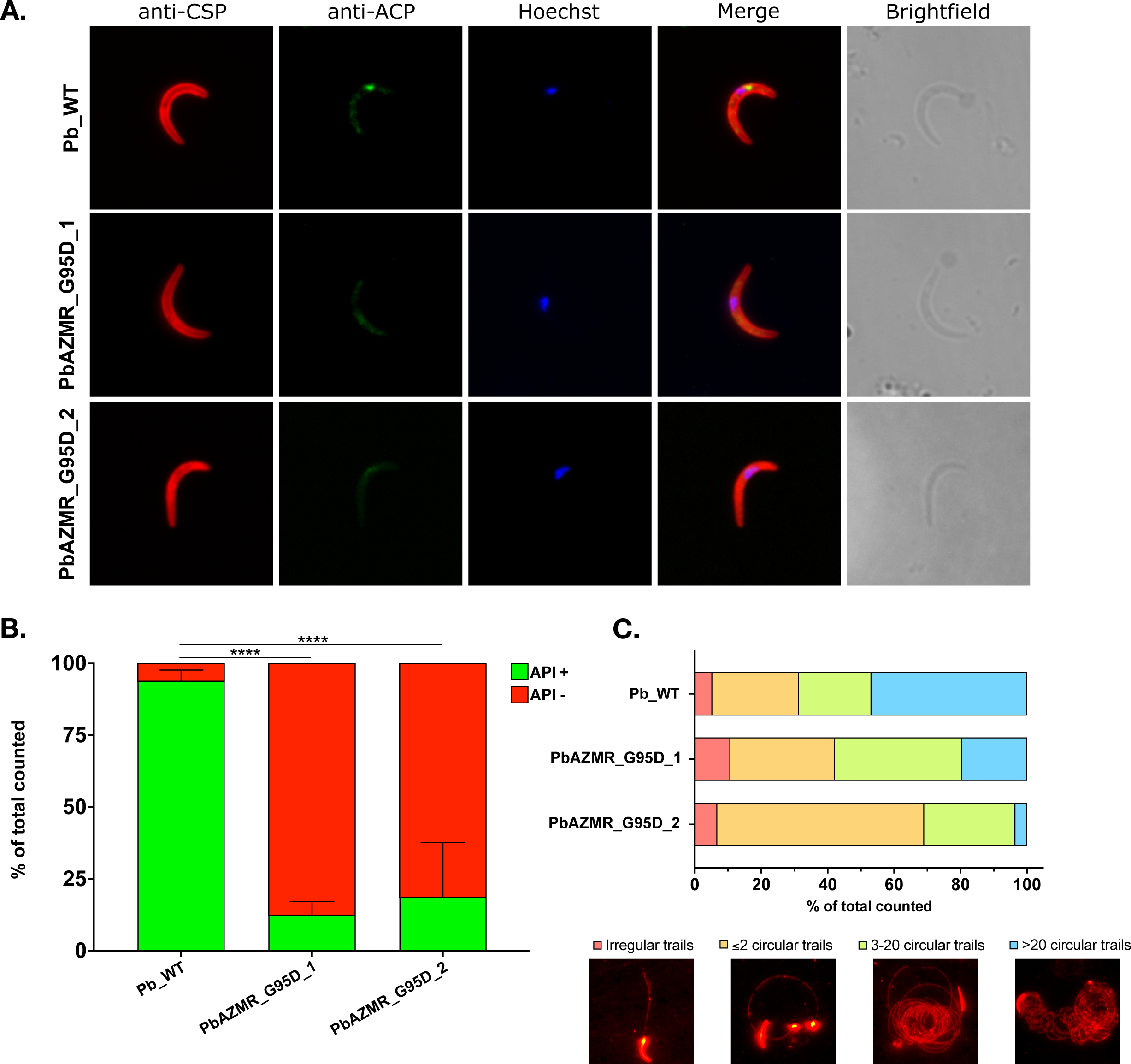
Immunofluorescence assay on PbAZMR_G95D_1, PbAZMR_G95D_2 and WT sporozoites reveals that azithromycin resistant sporozoites lose their apicoplast. A.) Sporozoites were stained with anti-circumsporozoite protein (CSP; red) to mark the outer surface of the sporozoite, anti-ACP (green) to mark the apicoplast, and Hoechst (blue) to mark the nucleus. WT sporozoites have a clear, punctate apicoplast (top panel), while azithromycin resistant sporozoites have diffuse apicoplast marker staining (bottom two panels). Images are representative of the calculated majority; B.) Quantification of apicoplast presence (API+, green) and absence (API-, red) in WT (n=2068), PbAZMR_G95D_1 (n=445), and PbAZMR_G95D_2 (n=855) sporozoites, where n=number of total sporozoites counted. A minimum of 110 sporozoites were counted for each parasite strain, for each of three independent mosquito infections. Green bars indicate the mean percentage apicoplast positive apicoplast, error bar represents +95% CI. Both azithromycin resistant strains produce sporozoites with significantly fewer intact apicoplasts than WT (P<0.0001, Fisher’s exact test); C.) Quantification of sporozoite motility trails from PbAZMR_G95D_1 (n=159), PbAZMR_G95D_2 (169), and WT (n=175) sporozoites, from two independent mosquito infections. There was no significant difference in types of trails observed between each parasite line (chi-square test).

To assess potential viability of these azithromycin resistant sporozoites, we performed a sporozoite gliding motility assay comparing the movement of PbAZMR_G95D_1 and PbAZMR_G95D_2 sporozoites to Pb_WT sporozoites (Figure 3C). PbAZMR_G95D_1 and PbAZMR_G95D_2 sporozoites are somewhat impaired in their gliding motility compared to Pb_WT, but the differences were not statistically significant (Figure 3C). Motility of the PbAZMR_G95D_1 and PbAZMR_G95D_2 sporozoites, despite mostly lacking a distinct apicoplast, is congruent with the fact that they had successfully made the journey from the midgut to the mosquito salivary glands (Figure 2B). Previous studies have noted, however, that even a modest reduction in sporozoite gliding can severely impact *in vivo* infectivity of sporozoites ^46, 47^. Thus, the motility differences observed here, although not statistically significant, may be biologically relevant to the infectivity of azithromycin resistant sporozoites, particularly *in vivo*.

### Azithromycin resistant P. berghei parasites are critically diminished in their capacity to establish an infection in a naïve vertebrate host

After assaying the *P. berghei* azithromycin resistant parasites through mosquito stage development and showing that they produce fewer sporozoites—most of which lack a distinct apicoplast—we next tested their ability to establish a patent infection in a naïve vertebrate host, either by bites from infected mosquitoes or direct intravenous (IV) injection of isolated sporozoites. Neither PbAZMR_G95D_1 or PbAZMR_G95D_2 was able to establish a patent infection in naïve mice bitten by at least 15 infected *A. stephensi* mosquitoes (Figure 4A). Similarly, IV injection of 1×10^3^ PbAZMR_G95D_1 or 1×10^3^ PbAZMR_G95D_2 sporozoites failed to produce patent infections (Figure 4B), whereas Pb_WT sporozoites reliably produced infections with a time-to-patency (as observed by microscopy of Giemsa stained thin blood smears) of 3-4 days (Figures 4A and B). When the inoculum of IV injected sporozoites was increased to 1×10^4^ sporozoites, PbAZMR_G95D_2 sporozoites were able to establish a patent blood stage infection in all of six attempts (Figure 4C). It is important to note that patency for the resistant mutant required 4-13 days (mean 8.5 days ± 3.8 days; Figure 4C), whereas Pb_WT sporozoites produced a patent infection in 3 days for all of 10 attempts (Figure 4C). DNA sequencing confirmed that the G95D Rpl4 mutation was present in the newly established PbAZMR_G95D_2 passage zero (P0) infection (data not shown). PbAZMR_G95D_1 sporozoites were unable to produce a patent infection in a naïve host in five attempts of IV injection of 1×10^4^ sporozoites (Figure 4C). Thus, through substantial mechanical intervention to provide a large sporozoite inoculum directly into a vein, we were able to achieve transmission of PbAZMR_G95D_2 but not PbAZMR_G95D_1. Even then, the PbAZMR_G95D_2 parasites showed a clear fitness defect over and above their liver stage delay, with blood stage parasites failing to increase substantially in parasitaemia when monitored over 14 days post sporozoite IV injection (Figure 4D).

**Figure 4.**
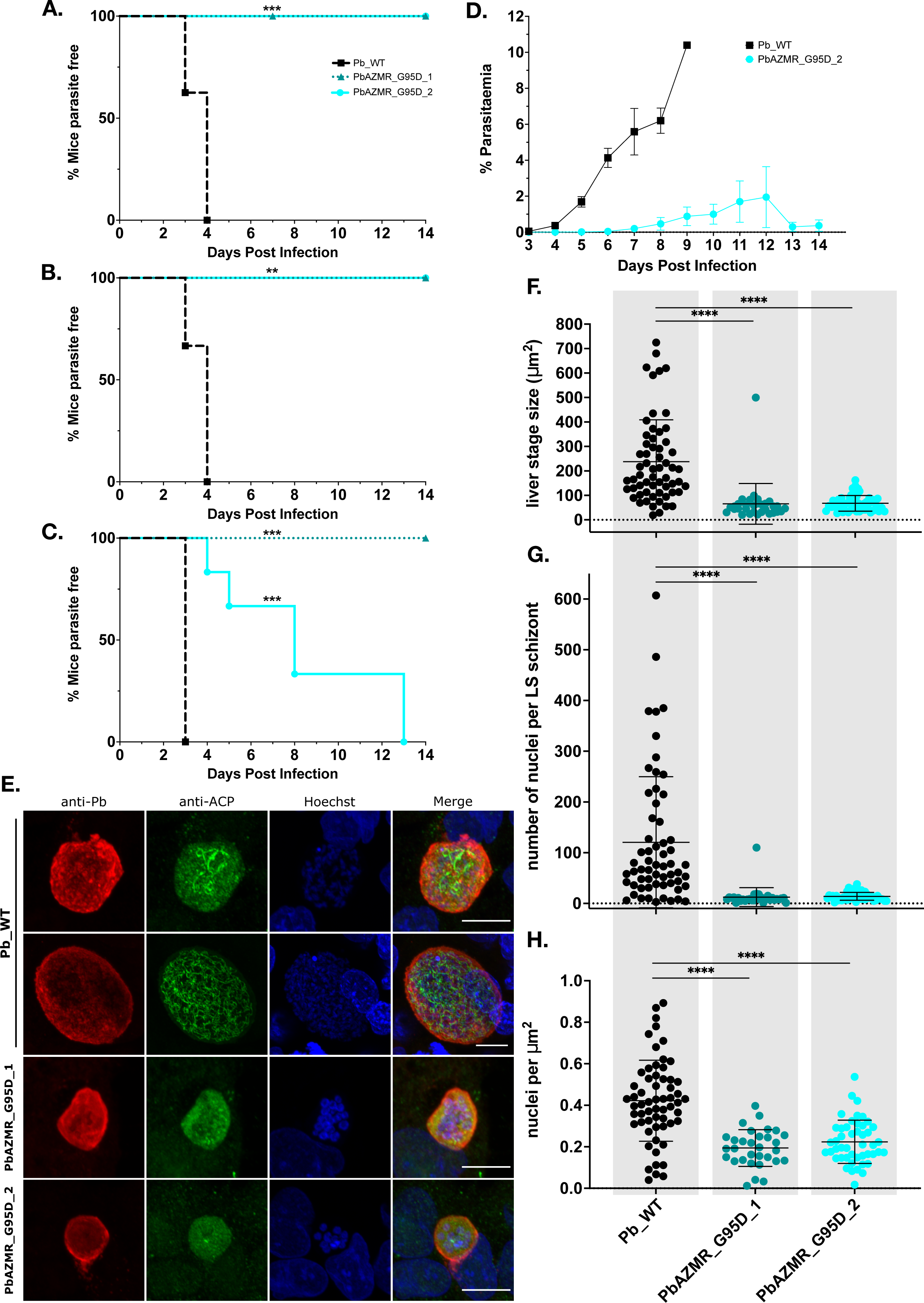
Transmission of azithromycin resistant *P. berghei* parasites to a naïve host is severely impaired, and *in vitro* liver stage development appears stalled. A.) Kaplan-Meier curves of P0 infections by biting. At least 15 infected mosquitoes were allowed to feed on naïve mice for at least 15 minutes and mice were monitored for 14 days by Giemsa smear (one mouse infected with PbAZMR_G95D_1 was only monitored until day 7, indicated by the initial triangle marker). Number of transmission attempts per strain were WT, n=8; PbAZMR_G95D_1, n=4; PbAZMR_G95D_2, n=4. Naïve mice infected with azithromycin strains by biting failed to establish an infection, and this was significantly different than WT (P=0.0007, Mantel-Cox test) B.) Kaplan-Meier curves of P0 infections by 1×10^3^ IV of sporozoites. Mice were monitored for 14 days by Giemsa smear. Number of transmission attempts per strain were WT, n=6; PbAZMR_G95D_1, n=1; PbAZMR_G95D_2, n=4. Naïve mice infected with azithromycin strains by 1×10^3^ IV failed to establish an infection, and this was significantly different than WT (P=0.01, Mantel-Cox test). C.) Kaplan-Meier curves of P0 infections by 1×10^4^ IV of sporozoites. Mice were monitored for 14 days by Giemsa smear. Number of transmission attempts per strain were WT, n=10; PbAZMR_G95D_1, n=5; PbAZMR_G95D_2, n=6. Only naïve mice infected with either WT or PbAZMR_G95D_2 by 1×10^4^ IV were able to establish an infection, but infection with PbAZMR_G95D_2 sporozoites showed delayed patency (mean 8.5 days ± 3.8 days [SD]) and impaired growth. This was significantly different than WT (P<0.0001, Mantel-Cox test). D.) Growth curve comparison of P0 infections established by WT and PbAZMR_G95D_2 by 1×10^4^ IV of sporozoites. Mice infected with PbAZMR_G95D_2 sporozoites grow slowly. Points shown indicate mean parasitaemia (±SEM), WT, n=9; PbAZMR_G95D_2, n=6, where ‘n’ represents the number of infected mice monitored for the period; E.) Immunofluorescence assay of liver cells infected with WT, PbAZMR_G95D_1 and PbAZMR_G95D_2 sporozoites in vitro and fixed at 48 hours post infection. The top two panels show HC-04 cells infected with WT sporozoites, the middle two panels show HC-04 cells infected with PbAZMR_G95D_1 sporozoites and the bottom two panels show HC-04 cells infected with PbAZMR_G95D_2 sporozoites. Azithromycin resistant liver stage schizonts only show dispersed ACP (apicoplast) staining. Nuclei (blue) in azithromycin resistant liver stage schizonts appear clustered and donut-shaped. Red staining indicates anti-Pb, marking the outside of the liver stage parasite. All images are maximum projections and scale bars represent 10 μm; F.) Quantification of parasite size (μm^2^) at 48-hours post sporozoite inoculation of HC-04 cells *in vitro* pooled from two independent experiments, with WT (n=60), PbAZMR_G95D_1 (n=31), and PbAZMR_G95D_2 (n=47), where ‘n’= number of individual liver stage schizonts measured. PbAZMR_G95D_1 and PbAZMR_G95D_2 liver stages are significantly smaller than WT (P<0.0001, Ordinary one-way ANOVA with Dunnett’s multiple comparisons test; bars represent the mean ±SD); G.) Pooled quantification of nuclei counts from the same liver stage schizonts quantified in F. PbAZMR_G95D_1 and PbAZMR_G95D_2 liver stages have significantly fewer nuclei than WT (P<0.0001, Ordinary one-way ANOVA with Dunnett’s multiple comparisons test; error bars represent the mean ±SD); H.) Nuclei counts, normalised to the size of each individual liver stage parasite described in F (number of nuclei per μm^2^). PbAZMR_G95D_1 and PbAZMR_G95D_2 liver stages have significantly fewer nuclei than WT even when normalised to liver stage parasite size (P<0.0001, Ordinary one-way ANOVA with Dunnett’s multiple comparisons test; error bars represent the mean ±SD).

These data again suggest there is a difference in phenotype penetrance between PbAZMR_G95D_1 and PbAZMR_G95D_2, which recapitulates similar differences during mosquito stage development. Despite both resistant mutants having the same azithromycin resistance conferring Rpl4 mutation, there remains a slight fitness advantage in the PbAZMR_G95D_2 parasites over the PbAZMR_G95D_1 parasites. There is, however, still a significant deficiency in the PbAZMR_G95D_2 sporozoites in that they are only transmission-competent when directly IV injected; and then only in massive numbers (Figures 4A-C).

The clear difficulty (or block in the case of PbAZMR_G95D_1) in the ability of our *P. berghei* azithromycin resistant parasites to establish a patent infection in a naïve host points to liver stage developmental defects, and the significant delay in patency of P0 infections induced by PbAZMR_G95D_2 sporozoites is consistent with this. This is not surprising given the importance of a proper functioning apicoplast during *P. berghei* liver stage development ^48–51^. Indeed, while some transmission of PbAZMR_G95D_2 is feasible, the parasites exhibit impaired development during mosquito stages, liver stages, and transmission to naïve hosts, identifying a clear fitness cost associated with the G95D Rpl4 mutation.

### Late liver stage development of azithromycin resistant P. berghei parasites is impeded

To investigate further the apparently aberrant liver stage development of our *P. berghei* azithromycin resistant strains, we inoculated *in vitro* grown HC-04 hepatocytes with sporozoites and observed the parasites by IFA to measure size and observe the morphology of the apicoplasts and nuclei. Firstly, we wanted to confirm whether PbAZMR_G95D_1 sporozoites are even able to infect liver cells given that they were unable to establish a blood stage infection in a naïve host by any method attempted. Secondly, while PbAZMR_G95D_2 sporozoites are clearly impeded in their ability to establish a blood stage infection in a naïve host, we wanted to examine the reasons for the apparent liver stage developmental delay and requirement for such massive sporozoite numbers to establish an infection in a naïve vertebrate host at all. Thirdly, we wanted to observe any associated aberrations in apicoplast morphology and visualise development of nuclei during liver stage development of PbAZMR_G95D_1 and PbAZMR_G95D_2.

Preliminary analyses revealed no differences in liver stage schizont size between azithromycin resistant parasites and Pb_WT at 24-hours post inoculation (data not shown). We therefore concentrated our efforts, and limited sporozoite supply, on a later liver stage timepoint of 48-hours.

IFA of liver cells inoculated with PbAZMR_G95D_1, PbAZMR_G95D_2 or Pb_WT sporozoites after 48 hours of development revealed marked difference in morphology (Figure 4E). WT liver stage schizonts, irrespective of size, typically showed reticulated ACP staining, indicative of normal apicoplast morphology at this point of liver stage development ^52^ (top two panels, Figure 4E). Furthermore, the nuclei of Pb_WT liver stage schizonts showed clear differentiation with each nucleus beginning to be associated with a developing merozoite (top two panels, Figure 4E). Contrast this with the PbAZMR_G95D_1 and PbAZMR_G95D_2 liver stage schizonts at 48-hours post infection where the ACP staining marking the apicoplast was cloudy and diffuse (bottom two panels, Figure 4E), reminiscent of what is seen in oocysts and sporozoites with these strains. Additionally, the nuclei of PbAZMR_G95D_1 and PbAZMR_G95D_2 developing liver stage schizonts appeared aggregated, forming unusual donut-shaped clusters (bottom two panels, Figure 4E). These images are typical of what we observed for 48-hour liver stages.

To quantify these observations, we used maximum projections of 48-hour post infection IFAs over two biological replicates and measured the size of each liver stage schizont, and we also quantified the nuclei in each developing parasite (Figure 4F and 4G). Both PbAZMR_G95D_1 (n=31) and PbAZMR_G95D_2 (n=47) liver stage parasites were significantly smaller than Pb_WT (n=60; Figure 4F) and contained fewer nuclei (Figure 4G). Moreover, this reduction in nuclei was independent of liver stage parasite size because normalising number-of-nuclei to the area of the liver stage parasite optical section still showed significantly fewer nuclei in PbAZMR_G95D_1 and PbAZMR_G95D_2 parasites compared to Pb_WT (Figure 4H). These late liver stage data essentially recapitulate the phenotype seen during development throughout mosquito stages; i.e. aberrant apicoplast morphology, and inability to properly differentiate nuclei. In sum, these defects in apicoplast integrity and nuclei biogenesis at both the oocyst and liver stages of the life cycle appear to severely hamper or even abrogate transmission of our *P. berghei* azithromycin resistant parasites.

### Phenotypes converge: liver stage development of azithromycin resistant P. falciparum parasites is impeded similar to P. berghei

To investigate further the effects of apicoplast-encoded Rpl4 G76V azithromycin resistance mutation on the life cycle progression of *P. falciparum*, we measured the rates of liver cell traversal (3-hours post inoculation) and invasion (18-hours post inoculation) of Pf_AZMR and Pf_WT sporozoites (Figure 5A, 5B). Reflecting the apparently near-normal motility of azithromycin resistant *P. berghei* sporozoites, azithromycin resistant Pf_AZMR sporozoites were able to traverse (Figure 5A) and invade (Figure 5B) *in vitro* cultured HC-04 cells with comparable efficacy to Pf_WT sporozoites. Moreover, Pf_AZMR sporozoites harboured intact apicoplasts (data not shown), unlike the *P. berghei* azithromycin resistant parasites, which mostly lacked a distinct apicoplast (Figure 3B).

**Figure 5.**
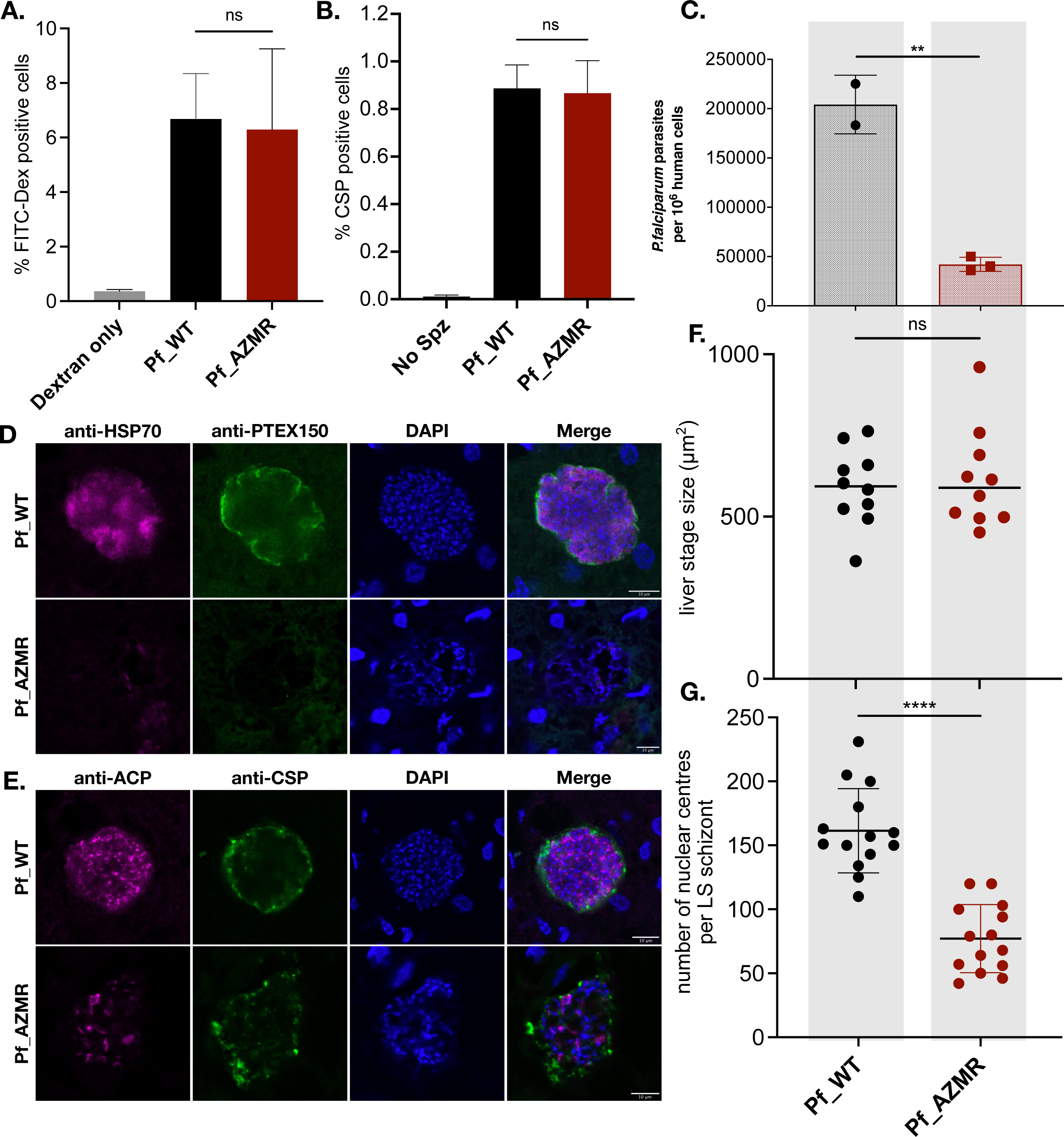
Liver stage development of *P. falciparum* azithromycin resistant parasites is impaired. A.) HC-04 cell traversal by Pf_WT and Pf_AZMR sporozoites at multiplicity of infection (MOI) 0.3, measured by FITC-Dextran uptake and counted by FACS. Cells were fixed at 3-hours post inoculation and show no significant difference between Pf_WT and Pf_AZMR indicating normal sporozoite motility in azithromycin resistant parasites (ns = not significant, Paired t-test two-tailed P value); B.) HC-04 cell invasion by Pf_WT and Pf_AZMR sporozoites measured by FACS after incubation for 18-hours using CSP-positive antibody staining of fixed permeabilized cells. No significant difference was seen in invasion of HC-04 between Pf_WT and Pf_AZMR sporozoites (ns = not significant, Paired t-test two-tailed P value); C.) Quantification of parasite liver load in humanised mice infected by i.v. injection of 8 x 10^5^ Pf_WT sporozoites (n=2 mice) or Pf_AZMR sporozoites (n=3 mice) by qRT-PCR. Livers of infected humanised mice were harvested at day 5 post sporozoite i.v. and tissue homogenised for subsequent loads analysis and determination of the degree of chimerism. Pf_AZMR infected mice showed a significantly lower liver stage parasite load than mice infected with Pf_WT sporozoites (Unpaired t-test, two-tailed P value, P=0.0023, error bars represent mean ±SD); D.) Immunofluorescence assay of liver sections from livers of humanised mice infected with either Pf_WT or Pf_AZMR sporozoites. Liver stage schizonts were stained with anti-HSP70 (cytosolic marker) and anti-PTEX150 (parasitophorous vacuole marker) and DAPI DNA stain. Pf_AZMR liver stage schizonts show little to no staining with either anti-HSP70 or anti-PTEX150 and show aberrant nuclear staining, with individual nuclei unclear. Scale bar 10 μm; E.) Immunofluorescence assay of liver stage schizonts and stained with anti-ACP (apicoplast marker), anti-CSP (parasite marker) and DAPI DNA stain. Pf_AZMR livers stage schizonts show a loss of anti-ACP and anti-CSP stain compared to WT, and again show aberrant nuclear staining, with DAPI appearing to aggregate around holes. Scale bar 10 μm; F.) Quantification of liver stage schizont size (μm^2^) from liver sections from livers of humanised mice infected with either Pf_WT or Pf_AZMR sporozoites. Sections from independent mice were selected and individual ‘n’ liver stage schizonts measured, Pf_WT (n=10), Pf_AZMR (n=10). No significant difference in liver stage schizont size was seen (ns = not significant, Unpaired t-test, two-tailed P value); G.) Quantification of nuclear centres counted from liver stage schizonts imaged in D and E. Pf_AZMR liver stage schizonts show significantly fewer nuclei than Pf_WT, indicating impaired liver stage development (Unpaired t-test, two-tailed P value, P<0.0001; error bars represent mean ±SD).

To mimic the *in vivo* life cycle phenotyping performed with azithromycin resistant *P. berghei* parasites, we infected humanised mice—with chimeric livers containing mostly human and some mouse hepatocytes—with Pf_AZMR sporozoites. Livers from humanised mice infected with either Pf_WT (n=two mice) or Pf_AZMR (n=three mice) sporozoites were harvested at day 5 post sporozoite IV inoculation and analysed by qRT-PCR to determine i/ the level of liver chimerism, and ii/ parasite loads. All mice had >75% human hepatocytes (data not shown), indicating a uniform infection potential, yet there was an 80% reduction in parasite load of Pf_AZMR parasites compared to Pf_WT (Figure 5C). Thus, Pf_AZMR parasites have a significantly reduced capacity to replicate in liver cells *in vivo*.

IFA on thin sections from these infected humanised mouse livers revealed defects in liver stage schizont development of Pf_AZMR in comparison to Pf_WT parasites (Figure 5D, 5E). The Pf_AZMR liver stage schizonts show little to no staining with either the cytosolic marker anti-HSP70 or the parasitophorous vacuole marker anti-PTEX150 (Figure 5D), indicative of severe overall morphological anomalies. Additionally, Pf_AZMR liver stage schizonts exhibit aberrant nuclear staining with no clear individual nuclei (Figure 5D). Indeed, nuclei often appeared to be aggregated (Figure 5E), reminiscent of the donut-shaped clusters of nuclei in *P. berghei* azithromycin resistant liver stages (Figure 4E). Moreover, whilst the size of the liver stage parasites was the same for Pf_AZMR in comparison to Pf_WT (Figure 5F), the azithromycin resistant parasites had significantly fewer nuclei (Figure 5G). Apicoplast morphology is also aberrant in Pf_AZMR late liver stage schizonts, with anti-ACP staining of Pf_AZMR revealing their apicoplasts to be collapsed and poorly defined in comparison to Pf_WT (Figure 5E).

Based on these severe defects of gross morphology, nuclear proliferation/separation, and apicoplast integrity for the Pf_AZMR parasites at liver stage, they appear to be experiencing severe fitness costs at day 5 of liver stage development attributable to the G76V Rpl4 mutation.

## DISCUSSION

### Rpl4 is a hotspot for macrolide resistance

Our four azithromycin selection regimens, across two *Plasmodium* species, all retrieved parasites with point mutations in the apicoplast-encoded ribosomal protein Rpl4. Mutations in Rpl4 are also associated with azithromycin resistance in other strains of *P. falciparum* ^30, 33, 43^, zoonotic infections of people with *Babesia microti* ^53^, in the plastid of the green alga *Chlamydomonas reinhardtii* ^54^, and in various bacteria related to the ancestor of plastids ^55, 56^. Mutations in Rpl4 thus seem to be a universal mechanism of resistance to azithromycin and closely related macrolides.

Why are apicoplast Rpl4 mutations so prevalent in malaria parasite azithromycin resistance? We previously argued that malaria parasite organelle genomes, which are multi copy ^57^, are hotspots for resistance generation simply because more chances exist for mutations to arise than do in single copy nuclear genes ^14^. Furthermore, organelle genomes could be heteroplasmic, allowing selection of pre-existing mutations by a drug to rapidly generate resistance, as likely occurs with atovaquone ^14^. Moreover, several different Rpl4 mutations can confer azithromycin resistance (Figure 1F), somewhat akin to different mutations in mitochondrial cytochrome *b* being able to confer atovaquone resistance ^58, 59^, which provides higher mathematical probability of resistance arising in a particular gene. Whatever the case, the frequency of Rpl4 mutations in resistance to azithromycin, both in this and other studies, demonstrates that apicoplast Rpl4 is a hotspot for azithromycin resistance. Sequencing Rpl4 should be a first point-of-call for surveillance of malaria parasite azithromycin resistance.

### Rpl4 mutations confer fitness advantage to parasites under drug pressure during the asexual, blood stages, but fitness deficits for transmission

Drug pressure is intermittent across the malaria parasite life cycle. Most drugs target the asexual blood phase in the human vertebrate host, yet drug pressure subsides when parasites progress through their sexual cycle in the mosquito host. Pressure can recommence if the parasites make it back into a new human host who is using antimalarials. At this point, a resistance-conferring mutation becomes an obvious selective advantage again, but the resistance allele(s) must transit the mosquito stages and the human liver to achieve transmission (and ultimately spread) of resistance. The fitness landscape for a parasite gene thus changes quite dramatically across the parasite life cycle.

Both *P. berghei* and *P. falciparum* are dependent on their apicoplast during mosquito stage development ^15–17, 60, 61^, and we set out to determine if mutations in apicoplast genes impacted parasite fitness in mosquitoes. Here, we showed that mutations conferring resistance to azithromycin (selected for by drug pressure during the blood phase) have differing consequences on fitness during mosquito phases. In *P. berghei,* the fitness deficit is severe in mosquito stages such as oocyst development and sporozoite production. However, in *P. falciparum* a different mutation in the same gene created no detectable fitness cost to the human parasite in mosquitoes. The difference between *P. berghei* and *P. falciparum* azithromycin resistant parasites in mosquito stage fitness likely relates to a combination of interacting factors. Firstly, the specific, resistance causing mutations in Rpl4 differ between parsite species (G95D or S89L in *P. berghei* versus G76V in *P. falciparum*; Figure 1E). Secondly, each parasite species is differentially reliant on its apicoplast in mosquitoes. For instance, based on the essentiality of apicoplast fatty acid biosynthesis for mosquito phase development in *P. falciparum* ^15^ but not rodent malaria species ^48, 49, 62^, one might expect *P. berghei* with a sub-optimal apicoplast to cope better in mosquitoes. However, that was not the case for *P. berghei* Rpl4 mutants phenotyped here, all of which exhibited marked reduction in fitness in mosquitoes. Differences in the mosquito host-parasite interaction could also impact the metabolic requirements for sporozoite development. *A. stephensi* is less susceptible to *P. berghei* than the natural mosquito host, *A. dureni* ^63^ whereas *A. stephensi* is an efficient natural host of *P. falciparum* ^64^. Thus, the differences in the infection rates by different parasites might reflect the evolutionary adaptation of each parasite to our provided mosquito host, as has been observed with immune responses in different mosquito/parasite interactions ^65, 66^.

If a mutant parasite is able to complete the mosquito lifecycle stages, and sporozoites accumulate in the mosquito salivary glands, its next phase—after being delivered by a bite—is to find and infect the vertebrate liver. Again, drug resistance mutations undergo selection for sufficient fitness during this liver phase. Interestingly, there is a convergence of phenotypes between *P. berghei* and *P. falciparum* azithromycin resistant parasites at late liver stage, both having defective apicoplasts and reduced/abnormal nuclei biogenesis. The malaria parasite liver stage is tremendously proliferative with extremely high metabolic demands ^67^, and our phenotyping indicates that a putatively sub-optimal apicoplast translation apparatus is apparently a major impediment to liver development of both *P. berghei* and *P. falciparum*, *in vitro* and *in vivo*.

In *P. berghei*, the combined burden of poor/abnormal growth of azithromycin resistant parasites in mosquitoes and liver blocked transmission, except where we provided massive mechanical intervention. In *P. falciparum*, a severe detrimental effect in the liver stage *in vivo* (in humanised mice) predicts a severe impairment of transmission. Whilst we were unable to test whether the Pf_AZMR parasites are competent to progress from liver stage to blood stage and thus achieve transmission, it seems likely that the *P. falciparum* azithromycin resistant mutant—despite performing better in mosquitoes—will nevertheless experience impediments to transmission. Whether or not the G76V mutation will block or reduce transmission of azithromycin resistance in a real-world setting is not yet possible to say, but the indications are that the mutation will not circulate freely, which should ultimately result in reduced transmission of azithromycin resistance, which gives azithromycin one more desirable characteristic as a potential partner drug. Refractoriness to spread of resistance would also mean that use of azithromycin for prophylaxis might pose less of a risk for resistance selection than is commonly assumed.

## ACKNOWLEDGEMENTS

Financial support from the National Health and Medical Research Council (GNT2016391, GNT1162550, GNT1143974), the Australian Research Council (FL170100008, LE200100181), and an Australian Postgraduate Scholarship (HB) are gratefully acknowledged. We thank Anton Cozijnsen, Vanessa Mollard and Ryan Steel for technical support.

## MATERIALS & METHODS

### Experimental animals and parasites

Male Swiss Webster mice, between 3 and 12 weeks old, were used in all experiments. Animals were sourced from Monash Animal Research Platform. All animal experiments were in accordance to the Prevention of Cruelty to Animals Act 1986 and the Prevention of Cruelty to Animals Regulations 2008 and reviewed and were permitted by the Melbourne University Animal Ethics Committee (Ethics IDs 1613928.1, 1714169, 1914889.1). Animals were anaesthetised using ketamine/xylazine and euthanised via slow fill carbon dioxide followed by cervical dislocation. *Plasmodium berghei* ANKA strain parasites were used in all experiments both as a reference wild type (WT) line and to generate all subsequent drug resistant parasite lines. *P. falciparum* NF54 strain parasites were used to generate all *P. falciparum* drug resistant parasite lines. Adult female *Anopheles stephensi* mosquitoes (MR4) were used in all experiments and reared under standard insectary conditions. Adult mosquitoes were grown at 27°C in 80% humidity, with a light/dark cycle of 14:10 hours. Mosquito infections were performed in dark conditions, and infected mosquitoes were maintained on 10% (v/v) sucrose.

### Drugs

Azithromcyin was obtained from Sigma-Aldrich (Merck, NSW Australia), as pure pharmaceutical secondary standards. Azithromycin was provided as a dihydrate (CAS Number: 117772-70-0) and administered once per day (q.d.) for 4 days as a suspension in PEG400 (MP Biomedicals, Ohio, CAS Number 25322-68-3) via oral gavage (OG). Drug dilutions were made fresh for each dosing experiment from identical drug batches as determined by LOT numbers.

### Activity of drugs *in vivo*

To test for the curative dose and to confirm the efficacy and tolerance of each drug, an initial *in vivo* drug dosing assay was performed. Briefly, a donor mouse was pre-infected with *P. berghei* ANKA parasites via intraperitoneal (IP) injection of frozen stock. Once the donor mouse reached 5% parasitaemia, it was euthanised and infected blood collected via cardiac puncture. Infected blood was diluted in RPMI 1640 medium (Gibco, Cat no. 61870-036) + 10% foetal bovine serum (Gibco, Cat no. 10100147) to a concentration of 1×10^7^ infected red blood cells (iRBC) per 200 μL. Two cohorts of three mice (placebo/vehicle treated [n=3] and drug treated [n=3]) were infected with 1×10^7^ iRBC via intravenous (IV) tail vein injection on day 0. Once mice reached 2% parasitaemia (approximately day 3), they were treated with the placebo/vehicle or drug by OG or IP injection (on days 3-6) and monitored daily by Giemsa (10% [v/v]) stained thin blood smears to determine parasite clearance. Mice were further monitored for health by daily weight measurements.

To evaluate the reported schizontocidal activity, dosing range and dosing methods of azithromycin, and to determine the 50% and 90% effective doses (ED50 and ED90 respectively) a Peters’ four-day suppressive test ^68^ was performed. Two donor mice were pre-infected with WT *P. berghei* ANKA parasites via IP injection of fresh blood from a P0 infection. Once the donor mice reached approximately 5% parasitaemia, they were euthanised and infected blood collected via cardiac puncture. Infected blood from both mice was mixed and diluted in RPMI 1640 medium + 10% foetal bovine serum to a concentration of 1×10^7^ iRBC per 200 μL. Seven cohorts of four mice were infected with 1×10^7^ iRBC via IV tail vein injection on day 0. All seven cohorts were administered the vehicle (200 μL PEG400) as a control or a given concentration of azithromycin (5 mg/kg, 15 mg/kg, 30 mg/kg, 45 mg/kg, 60 mg/kg, 70 mg/kg in 200 μL PEG400) q.d. by OG on days 0-3, with the initial dose on day 0 given 4 hours post-infection (pi). Parasitaemia was measured on day 4 by Giemsa-stained thin blood smears to determine percentage suppression in comparison to the untreated control. Mice were also monitored daily for health and to confirm parasites were being cleared. Percentage suppression (PS) of parasitaemia was calculated using the formula: PS=100 x ([C_AV_-T]/C_AV_), where C_AV_= average parasitaemia in the control group, and T=parasitaemia of each treated replicate. Each PS replicate was then plotted against the dose, and a non-linear regression curve was fitted using GraphPad Prism 9 (Version 9.1.0) to determine the ED50 and ED90 doses of azithromycin.

### Generating drug resistant *P. berghei* parasites

Selection of resistant *P. berghei* parasites *in vivo* was performed in the same way for each drug. After inoculation via IV injection with 1×10^7^ iRBCs from a WT *P. berghei* ANKA infected donor mouse, mice were treated with a clearance dose of the drug (60 mg/kg or 70 mg/kg azithromycin) administered as described above. Mice were monitored daily by Giemsa-stained thin blood smears, and once recrudescence was observed and parasitaemia reached approximately 1-2%, mice were treated with another clearance dose of the drug. Parasites were occasionally passaged into naïve mice should an immune response appear to have been suppressing growth. This recrudescence→treatment→recrudescence→treatment cycle was repeated until parasitaemia failed to reduce in the presence of the drug, at which point parasites were deemed resistant and parasite lines were cloned by limiting dilution. Cloned lines were then confirmed to be resistant to the clearance dose by performing an *in vivo* drug assay as described above.

### Generating drug resistant *P. falciparum* parasites

NF54 and 3D7 parasites were cloned by limiting dilution prior to resistance generation. Two independent clones of NF54 and one of 3D7 were used for selection. Parasite cultures containing 5×10^8^ parasites were treated continuously for 96 hours with 300 nM azithromycin or 72 hours with 30 nM atovaquone, or 200 nM pyrimethamine. All three parasite lines were also grown without drug as a sequencing control. Following treatment, cultures were allowed to recover in drug free media until recrudescence (28 days post azithromycin treatment, 30-38 days post atovaquone treatment, and 24-29 days post pyrimethamine treatment). Drug was reapplied to recrudescent parasites at the initial concentration. For azithromycin resistance, parasites grew normally at 300 nM azithromycin and continued to grow at concentrations up to 1.2 µM. Atovaquone and pyrimethamine resistant parasites required from two to five cycles of treatment and recovery before stable drug resistance at three times the initial concentrations was obtained. Resistance parasites were cloned by limiting dilution, and drug resistance confirmed by standard SYBRgreen drug assays. DNA was prepared from independent clones for whole genome sequening.

### Genomic DNA preparation and initial sequence analysis

Whole blood was collected via cardiac puncture from mice infected with clonal drug resistant *P. berghei* parasite lines, and 10 μL aliquots were transferred into 190 μL dPBS for a total volume of 200 μL. Genomic DNA was extracted using an Isolate II Genomic DNA Kit (Bioline, Meridian Bioscience Australia), following the manufacturer’s instructions. Known loci potentially involved in resistance to azithromycin amplified by polymerase chain reaction (PCR) for targeted sequencing (See Supplementary Table 1. for primer sequences and PCR conditions). Correct sized amplicons were then purified using an Isolate II PCR and Gel Kit (Bioline, Meridian Biosciences, Australia) following the manufacturer’s instructions for PCR clean-up and sent for Sanger sequencing at the Victorian Clinical Genetics Services (VCGS; Murdoch Children’s Research Institute, Royal Children’s Hospital, Parkville, Victoria) following their purified DNA criteria. Alignment and analysis of sequenced amplicons was done using Sequencher (Gene Codes Corporation, USA).

### Sample preparation and genomic DNA extraction for Illumina sequencing

To obtain a concentrated, pure parasite sample, free of contaminant mouse leucocytes, platelets and other material, we performed a *P. berghei* schizont preparation. Whole blood was collected via cardiac puncture from mice infected with the clonal drug resistant *P. berghei* parasite lines, and placed into a 50 mL overnight culture in complete schizont culture medium (70% [v/v] RPMI 1640 medium, 30% [v/v] FBS and gentamycin (Gibco) 4×10^-2^ mg/mL). Approximately 20 hours post culture, schizonts were purified using a CS column (Miltenyi Biotec, Germany) assembled in a VarioMACS magnetic cell separator (Miltenyi Biotec, Germany) and eluted in RPMI 1640 medium. Purified schizonts were spun down in a swing-bucket centrifuge at 1500 RPM for 10 minutes, resuspended in 1 mL of dPBS, centrifuged again at 1500 RPM for 10 minutes and resuspended in 1 mL dPBS. Each 1 mL washed schizont preparation was then distributed into five 200 μL aliquots and genomic DNA was extracted immediately using a QIAmp DNA Blood Mini Kit (Qiagen) following the manufacturer’s instructions with one exception. After lysis at 56°C, preparations were centrifuged at maximum (13000 RPM) and supernatant was placed into a new 1.5 mL tube to remove digested haemoglobin. After this step, the protocol was followed verbatim. Samples were sent to the VCGS (Murdoch Children’s Research Institute, Royal Children’s Hospital, Parkville, Victoria) for Illumina Nextera XT library preparation and whole genome sequencing on either an Illumina HiSeq4000 or NovaSeq2 (150 bp paired-end reads).

### Bioinformatic analysis

Analyses were performed using the High Performance Computing Services at The University of Melbourne (Parkville, Victoria) or personal computers. Raw FASTQ files were mapped to the *P. berghei* ANKA or *P. falciparum* 3D7 reference genome (Version 43 (*P. berghei*), Version 44 (*P. falciparum*), PlasmoDB ^69^) using the Burrows-Wheeler Aligner ^70^ and the quality checked with FastQC ^71^ and QualiMap ^72^. Mapped reads were visualised using Integrated Genomics Viewer (IGV, version 2.8.4 ^73^). Duplicate reads were marked using Picard tools (version 2.18.27) MarkDuplicates tool. Variant calling was performed following the Genome Analysis Toolkit (GATK)^74^ best practices pipeline ^75^, using the HaplotypeCaller (in GVCF mode) and GenotypeGVCFs tools. Raw variants were then annotated using SnpEff ^76^ and filtered using SnpSift ^77^ based on quality scores, read depth and genome position (i.e. intergenic versus coding regions). Further manual filtering and analysis was performed using GNU/Linux coreutils and Awk programming language ^78^ based on biological relevance.

### Infection of mosquitoes with *P. berghei*

Female *A. stephensi* mosquitoes were infected with clonal *P. berghei* drug resistant lines and WT *P. berghei* ANKA to investigate the fitness, phenotype and transmissibility of drug resistant parasites. Approximately 200 μL frozen blood stock was used to infect a donor mouse by IP injection. Once parasitaemia reached between 5-8% parasite readiness for infection was confirmed by checking for exflagellation of microgametes by microscopy. Donor mice were then anaesthetised, and individually placed on an infection cage containing 100 female *A. stephensi* mosquitoes, which were allowed to feed for up to 40 minutes in a 20°C incubator at 80% humidity. The donor mouse was euthanised post feed. Infected mosquitoes were maintained on 10% (v/v) sucrose for up to 28 days, or when the experiment was ended.

### Oocyst quantification and immunofluorescence assays

Whole midguts were dissected from infected mosquitoes between 9- and 14-days post blood feed and either stained with 0.2% (v/v) mercurochrome to assess oocyst numbers, or prepared for immunofluorescence assay. In 24-well plates, whole midguts were dissected in dPBS and then fixed in 4% (v/v) paraformaldehyde in dPBS for 1 hour at room temperature. After washing in 1 mL fresh dPBS three times for 5 minutes on an orbital shaker, midguts were blocked and permeabilised overnight at 4°C in midgut blocking buffer (0.25% [v/v] Triton X-100, 5% [w/v] BSA [Sigma-Aldrich, Cat no. A4503-50G] in dPBS). Midguts were incubated with primary antibodies (apicoplast marker acyl carrier protein [ACP] anti-serum ^79^ at 1:250) diluted in midgut blocking buffer for a minimum of 4 hours at room temperature on an orbital shaker. After washing three times in 1 mL dPBS, midguts were incubated at room temperature for 2 hours on an orbital shaker with secondary antibodies (goat anti-rabbit IgG Alexa Fluor 488 [Abcam] at 1:4000) diluted in midgut blocking buffer. Midguts were then washed three times for 5 minutes in 1 mL dPBS on an orbital shaker, with the second wash containing Hoechst 33342 diluted to 1:10 000 and mounted on glass slides in Dako fluorescence mounting medium.

Immunofluorescence assays were imaged on a Nikon C2 confocal microscope and images assembled in ImageJ Fiji (Version 2.1.0/1.53c) ^80^.

### Quantitative PCR for organellar genome detection

Primers designed to target genes encoded in the nuclear, apicoplast and mitochondrial genomes (See Supplementary Table 1. for primer sequences and qPCR conditions) were used to perform quantitative PCR (qPCR) on oocyst stage parasites isolated as described above. All primers were first tested to ensure similar amplification efficiency under the assay conditions by generating standard curves (Supplementary Figure 1.). Genomic DNA was extracted from parasite lines using an Isolate II Genomic DNA Kit (Bioline, Meridian Bioscience Australia), following the manufacturer’s instructions, with one exception. Whole midguts were first dissected into 180 μL lysis buffer GL (Bioline) and 20 μL Proteinase K and incubated at 56°C for 1 hour to ensure efficient lysis of oocysts, the original protocol was then followed verbatim. All qPCR assays were performed on a CFX Connect Real-Time PCR Detection System (Bio-Rad). Genome copy number was calculated using the ΔΔC_T_ method ^81^, first normalising mean C_T_ values of apicoplast and mitochondrial genes within samples to the mean C_T_ of the nuclear genes, then using the WT *P. berghei* ANKA samples as a calibrator for the drug resistant lines.

### Sporozoite quantification and immunofluorescence motility assays

Eight-well chamber slides (Merck Millipore Cat no. PEZGS0816) were coated with either mouse-derived monoclonal circumsporozoite protein (CSP) anti-serum (1:1000) ^82^ or mouse-derived polyclonal *P. berghei* anti-serum (1:1000; made in our laboratory) diluted in dPBS, and incubated at 37°C for 1 hour. Salivary glands were dissected in 1640 medium + 10% foetal bovine serum from infected mosquitoes between 20- and 24-days post blood feed, and purified sporozoites were counted. After removing the coating antibody solution from the chamber slide, up to 2×10^4^ sporozoites from each drug resistant *P. berghei* parasite line or WT *P. berghei* ANKA were added to individual wells and allowed to settle and glide for 1 hour at 37°C. The supernatant was then removed and parasites fixed with 150 μL of 4% (v/v) paraformaldehyde in dPBS for 20 minutes at room temperature. After washing gently with dPBS three times for 5 minutes, blocking buffer (0.2% [v/v] Triton X-100, 3% [w/v] BSA in dPBS) was added to each well and left to incubate for 20 minutes at room temperature. Primary antibodies were then added to each well (mouse anti-CSP [1:500], mouse anti-*P. berghei* [1:500] or rabbit anti-ACP [1:250] as described above) diluted in 3% (w/v) BSA in dPBS. Wells were again washed three times for 5 minutes with dPBS before incubating with secondary antibodies (goat anti-rabbit IgG Alexa Fluor 488 [Abcam]; goat anti-mouse IgG Alexa Fluor 568 [Abcam]) diluted in 3% (w/v) BSA (all 1:2000). Wells were then washed three times for 5 minutes in dPBS, with the second wash containing Hoechst 33342 diluted to 1:10000, and mounted with Dako fluorescence mounting medium. Sporozoites were then imaged on a Leica DM6000B compound fluorescence microscope and images assembled in ImageJ Fiji (Version 2.1.0/1.53c) ^80^.

### Mosquito infection and analysis of parasite development

Seven-day old female *Anopheles stephensi* mosquitoes (originally provided by M. Jacobs-Lorena, John Hopkins University) were allowed to feed on asynchronous gametocytes at 0.6% stage V gametocytemia using water jacketed glass membrane feeders. Mosquitos were then sugar starved for 48 hours to kill any that had not fed or males. Those that survived were given di-ionised water through cotton wicks and sugar cubes. For oocyst quantification; midguts were dissected from ethanol killed mosquitos 7 days post feed and stained with 0.1% mercurochrome. Salivary glands were dissected for mosquitos 16 days post feed, crushed with a pestle, and then filtered through glass wool. Sporozoites were counted using a hemocytometer.

### Infection of naïve mice with *P. berghei* sporozoites and analysis of infectivity

Naïve mice were infected with drug resistant or WT *P. berghei* ANKA sporozoites either by IV tail vein injection or by direct blood feeding. For IV infection, salivary glands were dissected from infected mosquitoes between days 20 and 24 post blood feed directly into RPMI 1640 medium + 10% (v/v) FBS. Sporozoites were then purified and counted prior to injecting either 1×10^3^ or 1×10^4^ sporozoites per mouse. For direct blood feed, infected mosquitoes were allowed to bite anaesthetised naïve mice. A total of 15 mosquitoes were allowed to blood feed for 15 minutes or until fed. Parasitaemia of mice was monitored by Giemsa-stained thin blood smears from day 2 post IV or mosquito bite to assess blood stage patency.

### Liver stage development analysis *in vitro*

To investigate the development of parasites through the liver stage, human hepatoma HepG2 (ATCC No. HB-8065) or HC-04 (ATCC MRA-975) cell lines were cultured at 37°C and 5% CO_2_ in a T-25 culture flask (Corning) in HepG2 medium (Advanced MEM minus L-Glutamine [Gibco] + 10% [v/v] HI-FBS [Gibco] + 1% [v/v] penicillin-streptomycin + 2 mM Glutamax [Gibco] and 0.1% [v/v] amphotericin B). Once the HepG2 or HC-04 cells were grown to the desired concentration, 1×10^5^ to 2×10^5^ cells were seeded onto coverslips in a 24-well plate in 1 mL HepG2 medium per well. The following day, sporozoites were dissected directly into HepG2 medium, purified and counted and up to 2×10^4^ sporozoites were added into each well and allowed to settle and invade for 2 hours at 37°C and 5% CO_2_. Media was changed after the 2-hour invasion process, and then twice daily thereafter until cells were fixed. Infected HepG2 or HC-04 cells were fixed on the coverslips for 20 minutes with 4% (v/v) paraformaldehyde in dPBS, at either 24-or 48-hours post sporozoite infection. Coverslips were washed with dPBS three times for 5 minutes in the 24-well plate before adding 1 mL ice-cold methanol for storage (no more than 3 weeks) at 20°C. After removing the methanol, coverslips were washed three times for 5 minutes with dPBS before cells were permeabilised with 0.1% (v/v) Triton X-100 in dPBS for 20 minutes at room temperature. Washing steps were repeated, and cells were then blocked for 30 minutes at room temperature with 3% (w/v) BSA in dPBS. Coverslips were incubated with primary antibodies (mouse anti-*P. berghei* [1:500] and rabbit anti-ACP [1:250]) and secondary antibodies (goat anti-rabbit IgG Alexa Fluor 488 [Abcam]; goat anti-mouse IgG Alexa Fluor 568 [Abcam]) in 3% BSA in dPBS for one hour at room temperature, washing three times for 5 minutes with dPBS in between each incubation. Following the probing with secondary antibodies, coverslips were washed three times for 5 minutes in dPBS, with the second wash containing Hoechst 33342 diluted to 1:10000 and mounted with Dako fluorescence mounting medium on glass slides and sealed with clear nail polish. Infected HepG2 and HC-04 liver cells were imaged on a Nikon C2 confocal microscope, and images assembled and measured in ImageJ Fiji (Version 2.1.0/1.53c) ^80^.

### Humanised mice processing

FRG humanised mice were obtained from Yecuris. An inoculum of 8×10^5^ *P. falciparum* control NF54 sporozoites and 8×10^5^ Azithromycin resistant sporozoites freshly isolated from mosquito salivary glands were injected into each humanised mouse (2 mice infected with NF54 and 3 mice with Azithromycin resistant lines).

Livers were obtained 5.5 days post infection from CO_2_-euthanized mice, and individual lobes were cut as described ^83^. Half of the lobes were embedded in OCT and frozen in liquid nitrogen for IFA analysis, while half was emulsified into a single-cell suspension and frozen and −80 for subsequent gDNA extraction.

For quantification of parasite load in the chimeric livers of each mouse; gDNA was isolated from the single-cell liver suspensions, and Taqman probe-based quantitative PCRs were performed as previously described ^83–86^ using primers that would specifically amplify the 18S gene in each parasite line 18S_fwd (5’-GTAATTGGAATGATAGGAATTTACAGGT-3’) and 18S_rev (5’-TCAACTACGAACGTTTTAACTGCAAC-3’). Human and mouse genomes were quantified using oligonucleotides specific for prostaglandin E receptor 2 (PTGER2) from each species, as described ^84^. All probes were labelled 5’ with the fluorophore 6-carboxy-fluorescein (FAM) and contain a double quencher that includes and internal ZEN quencher and a 3’ Iowa Black quencher from IDT.

The following probes were used:

18S: 5’FAM TGCCAGCAG/ZEN/CCGCGGTA-3IABkFQ

hPTGER2: FAM/TGCTGCTTC/ZEN/TCATTGTCTCG/3IABkFQ,

mPTGER2: FAM/CCTGCTGCT/ZEN/TATCGTGGCTG/3IABkFQ

Standard curves were prepared by titration of from a defined number of DNA copies for *P. falciparum* NF54, human and mouse controls. PCRs were performed on a Roche LC80 using LightCycler 480 Probes Master (Roche).

### Immunofluorescence assay

Purified NF54 and azithromycin resistant sporozoites were fixed with 4% paraformaldehyde (PFA)/ 0.0075% glutaraldehyde in PBS at room temperature for 20 minutes. Sporozoites were permeabilised in 0.1% Triton X 100 for 10 minutes before being blocked in 3% BSA PBS for 1 hour at room temperature. Samples were incubated with primary antibodies; rabbit anti-ACP antibody (1:500), and mouse anti-CSP 2A10 (1:1000), for 1 hour in 3% BSA PBS followed by secondary antibodies; anti-mouse 488 and anti-rabbit 594 (1:1000, Invitrogen). Images of NF54 or azithromycin sporozoites were taken at random based on CSP and DAPI and presence or absence of apicoplast was then determined by staining with ACP.

For liver stage immunofluorescence analysis; livers were perfused with 1x PBS and fixed in 4% PFA PBS overnight before being exchanged for sucrose and snap frozen in OCT (Invitrogen). Sections of 8 mm were cut on the cryostat apparatus HMM550. Samples were permeabilised using ice cold 10% methanol 90% acetone and blocked using 3% BSA PBS. Sections were then incubated with primary antibodies; mouse anti-CSP (2A10; 1:1000), mouse anti-HSP70 (1:500), rabbit anti-PTEX150 (1:500), mouse anti-EXP2 (1:500), rabbit anti-ACP (1:500) diluted in 3% BSA PBS. Secondary antibodies were goat anti-rabbit Alexa 594 and anti-mouse Alexa 488 (1:1000; Invitrogen). All samples were then incubated with DAPI (4’,6’-diamidino-2-phenylindole) at 1 ug/ml in PBS to visualize DNA and mounted in Prolong Gold mounting media (Invitrogen). Images were acquired on the Zeiss LSM 980 microscope.

Z stacks of both NF54 and azithromycin resistant liver stages were acquired using the same settings to allow quantitative analysis between different samples. The number of nuclear centres were counted by eye based on DAPI signal at the centre of each Z stack. For analysis of parasite size; a region of interest was drawn around each parasite based on DAPI signal and then the membrane area was calculated using FIJI software.

### Traversal assay

Cell traversal was measured using a cell wounding assay ^87^. HC-04 hepatocytes (1×10^5^) were seeded onto the bottom of a 96-well plate using IMDM (Life Technologies, 11360-070). Sporozoites were added at an MOI of 0.3 and left to traverse hepatocytes for 2.5 hours in the presence of 1 mg ml-1 FITC-labelled dextran (10,000 MW, Sigma Aldrich). Cells were trypsinized to obtain a single cell suspension for FACS analysis. For each condition, triplicate samples of 10,000 cells were counted by flow cytometry in each of the three independent experiments.

### Invasion assay

Assay was performed as previously described ^87^. Briefly, HC-04 hepatocytes (1×10^5^) were seeded onto the bottom of a 96-well plate using IMDM (Life Technologies, 11360-070). Sporozoites were added at an MOI of 0.3 and left to invade hepatocytes for either 5 or 18 hours before being trypsinized to obtain a single cell suspension and transferred to a 96 well round bottom plate. Cells were fixed and permeabilised (BD bioscience) and stained with Alexa fluor 647 conjugated mouse anti-CSP antibody in 3% BSA permeabilizing solution. Cells were then washed in PBS and analysed by Flow cytometry. For each condition, triplicate samples of 10,000 cells were counted in each of the three independent experiments.

---

**Figure S1.**
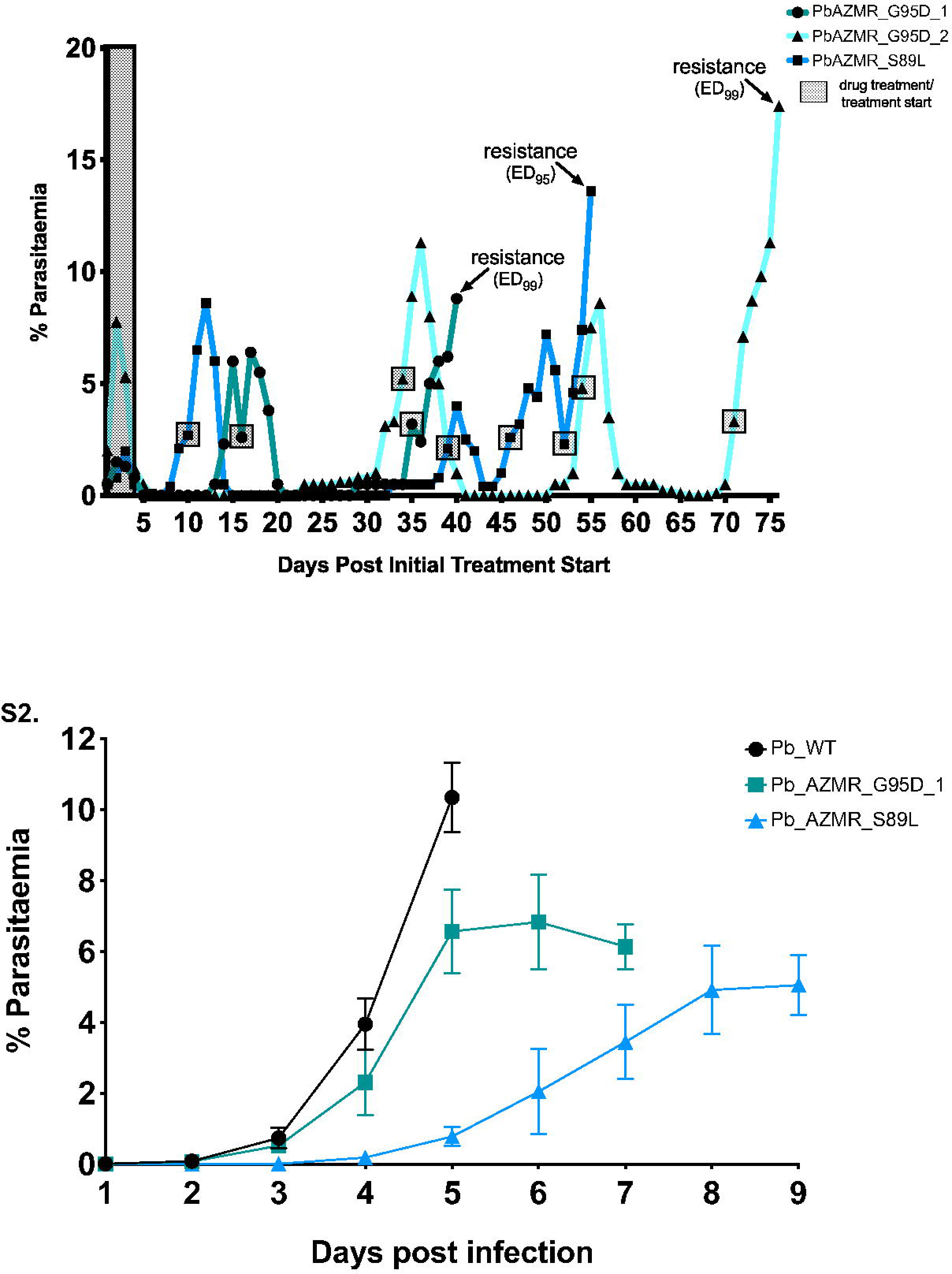
Generation of azithromycin resistant *P. berghei* parasites *in vivo*. PbAZMR_G95D_1 and PbAZMR_G95D_2 were both generated off a 70 mg/kg q.d. dosing regimen, and PbAZMR_S89L was generated off a 60 mg/kg q.d. dosing regimen. The initial four-day dose is represented by the shaded rectangle, and the start of each dose thereafter is indicated by a shaded box to avoid rectangle overlaps. Parasites were deemed resistant when they failed to reduce in parasitaemia on the third day post treatment start (with the exception of PbAZMR_S89L, which grew slowly so was monitored differently).

**Figure S2.** Blood stage growth of *P. berghei* azithromycin parasites is impaired compared to WT without drug pressure. Growth assay of PbAZMR_G95D_1 (n=3) and PbAZMR_S89L (n=3) compared to WT (n=3). Parasitaemia was monitored by Giemsa-stained thin blood smears. Dot points represent mean parasitaemia, and error bars indicate ±SD. Growth of both azithromycin resistant parasite lines is delayed compared to WT, and eventually fails to increase.

